# Cell non-autonomous functions of S100a4 drive fibrotic tendon healing

**DOI:** 10.1101/516088

**Authors:** Jessica E. Ackerman, Valentina Studentsova, Katherine T. Best, Emma Knapp, Alayna E. Loiselle

## Abstract

Identification of pro-regenerative approaches to improve tendon healing is of critical importance given the diminished quality of life and physical function that accompanies the typical fibrotic response to tendon injury. S100a4 modulates fibrosis through tissue-type dependent mechanisms, and the role of S100a4 in fibrotic, scar-mediated tendon healing has not been established. In the present study we tested the hypothesis that inhibition of S100a4 improves tendon function following acute injury and surgical repair. We demonstrate cell non-autonomous functions of S100a4 as S100a4 haploinsufficiency promotes regenerative tendon healing, including decreased scar formation and improved mechanical properties. Moreover, inhibition of S100a4 via antagonism of its putative receptor, the Receptor for Advanced Glycation Endproducts (RAGE), also decreases scar formation. Mechanistically, knock-down of S100a4 decreases myofibroblast and macrophage content at the site of injury, with both cell populations being key drivers of fibrotic progression. In contrast, S100a4^+^ cell depletion displays time-dependent effects on scar formation, and consistent impairments in restoration of mechanical properties, indicating a critical role for these cells in re-establishing tendon strength after injury. Finally, we demonstrate, that S100a4-lineage cells become α-SMA^+^ myofibroblasts, via loss of S100a4 expression. Using a combination of genetic mouse models, small molecule inhibitors and in vitro studies we have defined S100a4 as a novel, promising therapeutic candidate to improve tendon function after acute injury.

## Introduction

Tendons are composed primarily of a dense, highly aligned collagen extracellular matrix (ECM), and connect muscle to bone to transmit mechanical forces throughout the body. Following injury, tendon demonstrates limited regenerative potential and heal through a scar-mediated fibrotic process involving abundant, disorganized ECM deposition. While scar tissue can impart some mechanical strength to the healing tissue, it is also mechanically inferior to native tendon and dramatically impairs normal tendon function resulting in substantial morbidity. In addition, scar tissue increases tendon bulk and forms adhesions to the surrounding tissues, impeding normal range of motion (ROM). This pathological response to injury represents a major clinical burden considering there are over 300,000 surgical tendon repairs in the United States annually (1), and up to 40% of primary flexor tendon repairs heal with function limiting scar tissue (2). Despite this burden, there is currently no consensus biological or pharmacological approach to improve tendon healing, due in large part to a paucity of information on the cellular and molecular components involved.

S100a4 (also known as *Fsp1*, *Mts1*, *Pk9a*) is a member of the S100 family of EF-hand Ca^2+^-binding proteins, and is a potent regulator of fibrosis in many tissues including the liver (3, 4), lung (5), heart (6) and oral submucosa (7). An increase in the proportion of S100a4^+^ cells is characteristic of many fibrotic conditions (5, 8), and elevated serum S100a4 levels positively correlate with fibrosis clinically (4). Moreover, the therapeutic potential of S100a4 inhibition is suggested by S100a4-cell depletion studies and S100a4 RNAi treatments in which fibrosis was halted, or effectively reversed (4, 9, 10). While depletion of S100a4^+^ cells can inhibit fibrotic progression, S100a4 can also function as an intra- and extracellular signaling molecule to impact cellular processes including motility, survival, differentiation, and contractility (11, 12). Additionally, the effects of S100a4 are cell and tissue-type dependent. We have previously shown that *S100a4*-Cre efficiently targets resident tendon cells (13), however, the specific function of S100a4 and whether that function is cell-autonomous or cell-non-autonomous during scar-mediated fibrotic tendon healing is unknown.

In the present study we delineate the relative contributions of S100a4 expression and S100a4^+^ cells to scar-mediated tendon healing and investigated both the cell non-autonomous extracellular signaling function of S100a4, as well as the fate and function of S100a4-lineage cells. We have identified S100a4 haploinsufficiency as a novel model of regenerative tendon healing and define a requirement for S100a4^+^ cells in the restoration of mechanical properties during tendon healing. These data identify S100a4 as a novel target to improve tendon healing and demonstrate the efficacy of pharmacological inhibition of S100a4 signaling to improve functional outcomes during healing.

## Methods

### Ethics statement

All animal studies were approved by the University of Rochester Committee for Animal Resources.

### Mouse strains

S100A4-GFP^promoter^ mice (#012893), S100a4-TK (#012902), S100a4^GFP/+^ (#012904), S100a4-Cre (#012641), ROSA-Ai9 (#007909), and C57BL/6J (#000664) were acquired from The Jackson Laboratory (Bar Harbor, ME).

S100a4-GFP promoter mice contain a construct encoding EGFP under control of the S100a4 promoter sequence, resulting in green fluorescence in cells actively expressing S100a4 (14).

S100a4-TK mice contain a viral thymidine kinase gene downstream of the S100a4 promoter, and treatment with the nucleoside analog ganciclovir (GCV) halts DNA replication in proliferating cells expressing S100a4, resulting in apoptosis and S100a4^+^ cell ablation (9). Mice were treated twice per day (i.p) with 75mg/kg GCV. S100a4^GFP/+^ mice contain a GFP-encoding gene knocked into the exons 2-3 of the S100a4 gene, resulting in a 50% reduction in S100a4 protein expression (15). For RAGE Antagonist Peptide (RAP) studies, C57BL/6J mice were treated with either 100μg RAP or vehicle (0.5% bovine serum albumin in saline) via i.p. injection on D5-10 post-surgery.

### Murine model of tendon injury and repair

Male and female mice aged 10-12 weeks underwent complete transection and surgical repair of the flexor digitorum longus (FDL) tendon as previously described (16). Mice were monitored and given analgesics post-operatively as needed.

### RNA extraction and qPCR

For in vivo studies, the tendon repair site was excised from the hind paw at D10 following injury, along with 1-2mm of native tendon on either side. Three repairs were pooled, and RNA was extracted with TRIzol reagent (Life Technologies, Carlsbad CA). cDNA was generated with 500ng of RNA using an iScript cDNA synthesis kit (BioRad, Hercules CA). Quantitative PCR was carried out with gene specific primers, and expression normalized to *β-actin*. All experiments were done in triplicate and repeated twice.

### Assessment of gliding function and mechanical properties

Following sacrifice, the hindlimb was harvested at the knee. The medial side of the hindlimb was carefully dissected to free the FDL, and the proximal end was secured between two pieces of tape with cyanoacrylate. The distal tendon was loaded via the tape with weights ranging from 0 to 19g, with digital images taken upon application of each weight. MTP Flexion Angle and Gliding Resistance were calculated as previously described (13, 17, 18), with lower MTP Flexion Angle and higher Gliding Resistance corresponding to restricted range of motion and impaired gliding function. Following gliding testing, the FDL was released from the tarsal tunnel, and the proximal end of the tendon and the digits were secured in opposing custom grips on an Instron 8841 uniaxial testing system (Instron Corporation, Norwood, MA). The tendon was loaded until failure at a rate of 30mm/minute (17).

### Histology, Immunohistochemistry and Immunofluorescence

Following sacrifice, hind paws were dissected just above the ankle, and underwent routine processing for paraffin or frozen sectioning. Paraffin samples were fixed for 72h in 10% NBF, then decalcified for 2 weeks in 14% EDTA before processing. Three-micron sagittal sections were stained with Alcian Blue Hematoxylin / Orange G (ABHOG) or picrosirius red stain (Polysciences Inc, Warrington PA). Frozen samples were fixed overnight, decalcified for 4 days, incubated in 30% sucrose (in PBS) overnight, and embedded in Cryomatrix (#6769006, ThermoFisher, Waltham MA). Eight-micron sagittal sections on Cryofilm tape (Section-lab, Hiroshima, Japan) were cut on a Leica CM1860UV cryostat, and adhered to slides with 1% chitosan in 0.25% acetic acid.

Chromogen immunohistochemistry was performed on paraffin sections for S100a4 (1:20000, #197896, Abcam, Cambridge MA), with a rabbit polymer kit (MP-7401, Vector Laboratories, Burlingame CA). Immunofluorescence was carried out with the following primary and secondary antibodies: RAGE (1:100, #sc-365154, Santa Cruz Biotechnology, Dallas TX), with goat anti-mouse AlexaFluor488 secondary (1:1000, #A11029, ThermoFisher, Waltham MA); F4/80 (1:500, #sc-26643, Santa Cruz Biotechnology, Dallas TX) with a donkey anti-rabbit AlexaFluor594 secondary (1:200, #705-546-147, Jackson ImmunoResearch, West Grove PA); GFP (1:5000, #ab6673, Abcam, Cambridge MA), with a donkey anti-goat 488 secondary (1:200, #705-546-147, Jackson ImmunoResearch, West Grove PA); RFP (1:500, #ab62341, abcam, Cambridge MA), with a donkey anti-rabbit 647 secondary (1:200, #711-606-152, Jackson ImmunoResearch, West Grove PA); and α-SMA-Cy3 (1:250, #C6198 Sigma-Aldrich, St Louis MO). All slides were imaged with the Olympus slide scanner and processed with Olyvia software (Olympus, Waltham MA). Images were pseudo-colored using ImageJ software (v1.51j8, NIH). At least three animals were evaluated per genotype per time point.

### In Vitro Studies

Bone marrow derived primary macrophages (BMDM) were grown from the bone marrow of C57BL/6J mice. Following sacrifice, femurs were flushed with ice-cold phosphate buffered saline (PBS, without Ca^2+^ /Mg^2+^). The cell suspension was strained through a 70µm filter, resuspended in differentiation medium(19) and plated at a concentration of 3×10^6^ cells per 10cm plate. At D7 of differentiation, primary macrophages were re-plated as needed for experimental use.

#### Migration assay

Primary BMDMs were seeded at confluence in 96-well Oris 96-well plates (Platypus Technologies, Madison WI) with silicon stoppers inserted and incubated overnight. Plugs were removed, and cells washed once with PBS (with Ca^2+^ /Mg^2+^) prior to addition of treatment. Vehicle or recombinant S100a4 protein was added to wells at 20, 50, 200, 500, and 1000ng/mL in quadruplicate, and cells allowed to migrate for 24h. Following migration, cells were gently washed with PBS and stained with NucBlue Live ReadyProbe (Life Technologies, Carlsbad CA). A detection mask was affixed to the bottom of the plate to obscure cells that had not migrated, and the plate read on a Synergy 2 plate reader (Biotek, Winooski VT) at 360ex/480em, and fluorescence normalized to vehicle treated cells.

#### Polarization

Primary BMDMs were seeded into 6-well plates at 85% confluence overnight. Cells were treated with either S100a4-RP (20-1000ng/mL) or vehicle for 24h. Cells were lysed in Trizol (Life Technologies, Carlsbad CA), and RNA extracted using the RNeasy mini kit (Qiagen, Germantown MD). Quantitative PCR was performed for markers of M1 (*iNOS*, *TNFα*, *CD86*, *CD64*) and M2 (*CD206*, *Arg1*, *IL-1ra*, *CD163*) polarization, and data normalized to *β-actin* expression.

### Statistical analyses

Statistically significant differences between genotypes or treatments in in vitro and in vivo studies were assessed by unequal variance unpaired t-test, with the exception of the S100a4 qPCR time-course, BMDM migration and polarization experiments, which were analyzed by one-way ANOVA, followed by Tukey’s multiple comparisons test. All analyses were conducted using GraphPad Prism software (v8.0.0, La Jolla CA). p values ≤ 0.05 were considered significant, with the following conventions: * = p ≤ 0.05, ** = p ≤ 0.01, and *** = p ≤ 0.001.

## Results

### S100a4 is expressed by resident tendon cells and the S100a4^+^ population expands during healing

Spatial localization of S100a4 was examined before and after tendon repair surgery in S100a4-Cre; ROSA-Ai9 reporter mice to trace S100a4-lineage cells (S100a4^lin+^; Fig. 1A & B), and S100a4-GFP^promoter^ mice to identify cells actively expressing S100a4 (S100a4^active+^; Fig. 1C & D). Most resident tendon cells were S100a4^lin+^ in the uninjured tendon. Following tendon repair, the S100a4-lineage population expanded with S100a4^lin+^ cells in the native and bridging scar tissue at D7 and D14 post-surgery (Fig. 1B). Many resident tendon cells were actively expressing S100a4 at baseline, however many S100a4^−^ cells were also present (Fig. 1D). Following injury, the S100a4^active+^ population also expanded, with abundant S100a4^active+^ cells in the bridging scar tissue from D3 to D14, with a marked decrease by D28 (Fig. 1D). Notably, the expansion of the S100a4^active+^ population was also observed in the healing Achilles tendon (Supplemental Fig. 1), suggesting potential conservation of S100a4 function between tendons. Consistent with changes in spatial expression over time, *S100a4* mRNA expression increased from D3 to a peak at D10, followed by a progressive decline through D28 (Fig. 1E).

**Figure 1:**
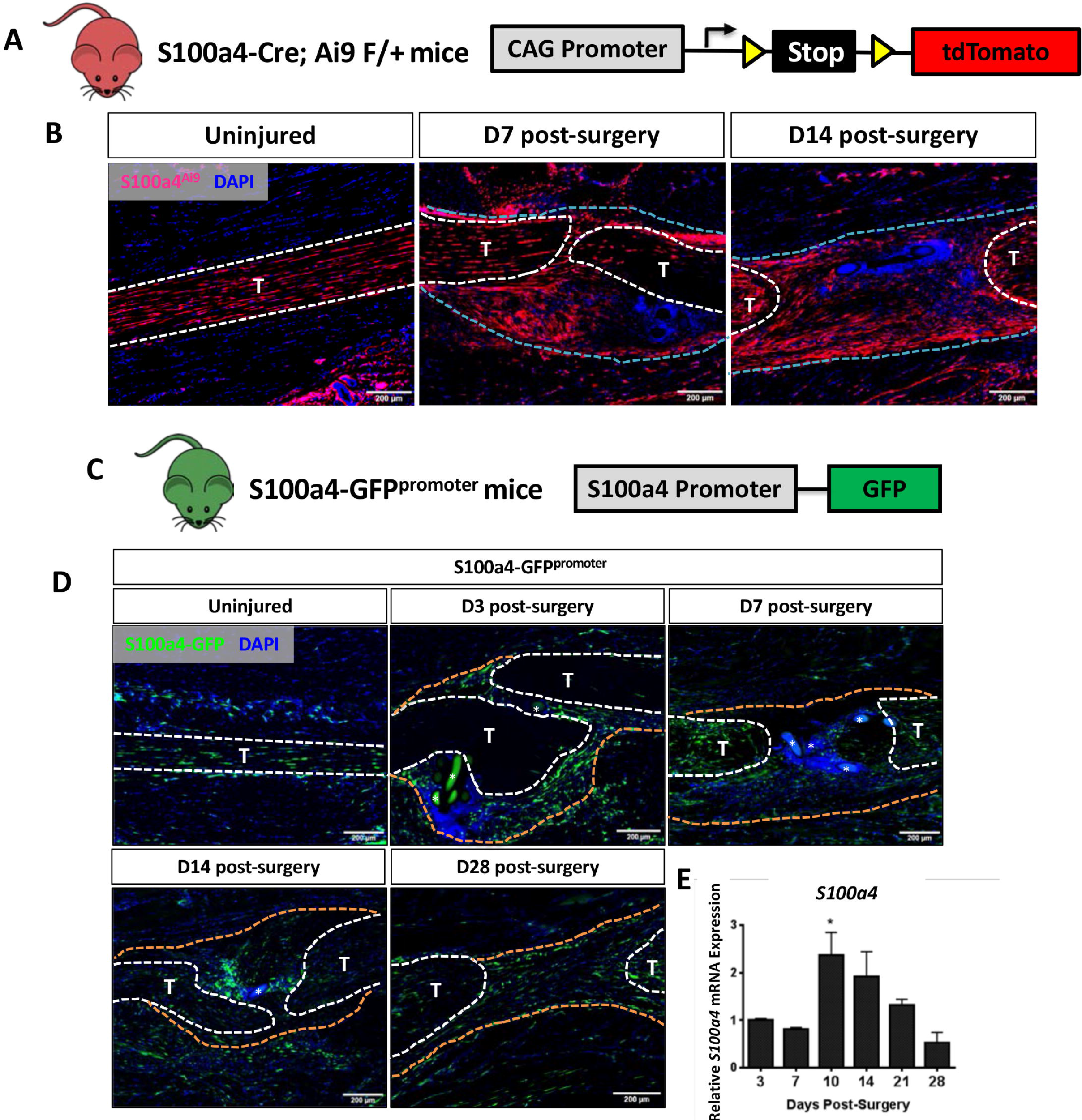
S100a4 is expressed by resident tenocytes and the S100a4^+^ cell population expands during tendon healing. (A & B) S100a4-Cre; Rosa-Ai9 reporter mice demonstrate efficient targeting of resident tendon cells. Following injury, the S100a4-lineage population expands, with S100a4^Lin+^ cells in the native tendon stubs and the bridging scar tissue at D7 and D14 post-surgery. Tendons are outlined in white, and bridging granulation tissue outlined in blue. (C) The S100a4-GFP^promoter^ construct identifies cells actively expressing S100a4 (S100a4^active+^). (D) A subpopulation of resident tenocytes is S100a4^active+^ at baseline, and the S100a4^active+^ population increases following injury, with S100a4^active+^ cells observed in the bridging scar tissue and native tendon ends through D28 post-surgery. Tendons are outlined in white, and bridging granulation tissue outlined in orange, (*) identifies sutures. (E) qPCR analysis of S100a4 during tendon healing demonstrates peak S100a4 expression at D10, followed by a progressive decline through D28 (n=3 per time-point). (*) indicates p<0.05 vs. D3 repair (1-way ANOVA). Data were normalized to expression in D3 repairs, and the internal control β-actin.

### S100a4 haploinsufficiency promotes regenerative, mechanically superior tendon healing

To determine the functional implications of decreasing S100a4 expression during tendon healing (Fig. 2A), we utilized S100a4 haploinsufficient mice (S100a4^GFP/+^), which results in a 50% reduction in *S100a4* mRNA expression in the tendon (Fig. 2B), as well as a robust decrease in S100a4 protein expression during tendon healing (Fig. 2C). Decreased S100a4 expression did not noticeably alter the spatial localization of S100a4^+^ cells in either the un-injured tendon or at D14 post-surgery (Supplemental Fig. 2). At D14 post-surgery, functional outcomes of scar formation in healing S100a4^GFP/+^ tendons were significantly improved compared to WT. A significant 36% increase in MTP Flexion Angle was observed in S100a4^GFP/+^ repairs, relative to WT (p=0.04) (Fig. 2D). Gliding Resistance was significantly decreased by 43% in S100a4^GFP/+^ repairs, relative to WT (p=0.028) (Fig. 2E), suggesting a reduction in scar formation in S100a4^GFP/+^ repairs. In addition, maximum load at failure was significantly increased (+35%) in S100a4^GFP/+^ repairs relative to WT (p=0.003) (Fig. 2F), while stiffness was increased 28% in S100a4^GFP/+^ repairs, relative to WT, however this increase was not statistically significant (p=0.08) (Fig. 2G). Taken together, these data suggest that S100a4 haploinsufficiency improves functional outcomes, while also improving tendon strength.

**Figure 2:**
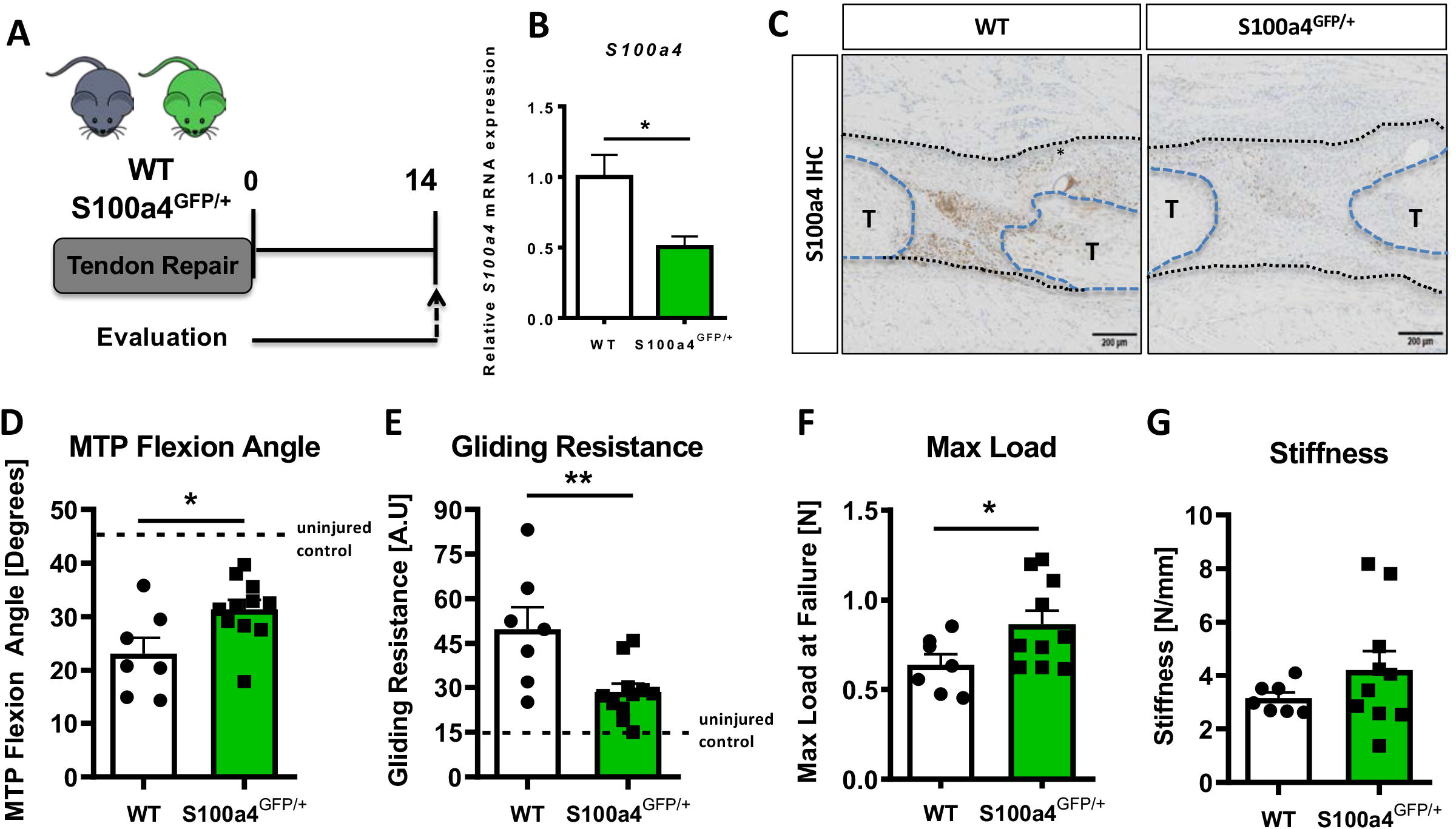
S100a4 knockdown promotes regenerative, mechanically superior tendon healing. (A) S100a4^GFP/+^ haploinsufficient and wild type (WT) littermates underwent transection and repair of the FDL tendon, and tendons were harvested at D14 post-surgery. (B) *S100a4* mRNA expression was reduced by 50% in S100a4^GFP/+^ tendon repairs, relative to WT (n=3 per group). (C) A substantial reduction in S100a4 protein expression was observed in S100a4^GFP/+^ tendon repairs, relative to WT. Tendon ends are outlined in blue and bridging scar tissue outlined in black (n=3-4 per group). (D-G) At D14, MTP Flexion Angle was significantly increased in S100a4^GFP/+^ (D), and Gliding Resistance was significantly decreased in S100a4^GFP/+^ repairs (E). Max load at failure was significantly improved in S100a4^GFP/+^ repairs (F), while no change in Stiffness was observed between genotypes (G) (n=7-10 per group). (*) indicates p<0.05, (**) indicates p<0.01 between genotypes, n=7-10 for (D-G) (un-paired t-test).

### S100a4 knockdown improves tendon morphology and decreases myofibroblast content

Morphologically, both ABHOG and picrosirius red staining demonstrate more highly aligned, mature collagen fibers bridging the tendon ends in S100a4 haploinsufficient mice (blue arrows, Fig. 3A). Consistent with this, *Col1a1* expression was significantly increased 5.6-fold (p=0.0095) in S100a4^GFP/+^ repairs, relative to WT repairs (Fig. 2B), while no change in *Col3a1* or *Scx* were observed between groups (Fig. 2C & D). Expression of the myofibroblast marker *α-SMA* was decreased 2.4-fold in S100a4^GFP/+^ repairs, relative to WT (p=0.02) (Fig. 2E). Consistent with this, a marked decrease in *α-SMA* protein was also observed in S100a4^GFP/+^ repairs, relative to WT (white arrows, Fig. 3F). These data suggest that S100a4 haploinsufficiency promotes regenerative tendon healing via deposition of a mature Col1 ECM and a decrease in pro-fibrotic myofibroblasts.

**Figure 3.**
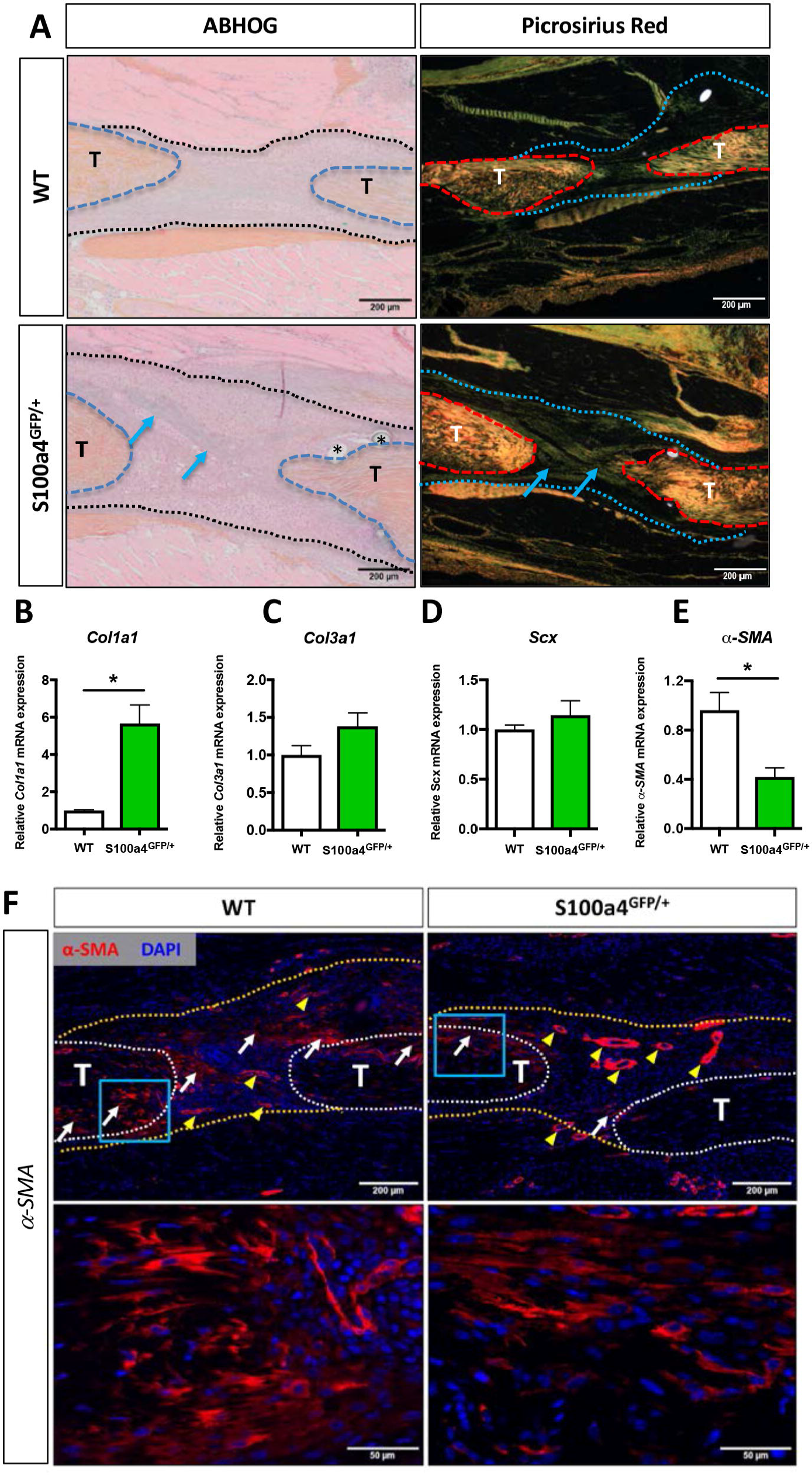
S100a4 haploinsufficiency enhances deposition of a mature Collagen matrix and reduced myofibroblast content. (A) ABHOG and picrosirius red staining demonstrate an increase in mature collagen fibers (blue arrows) bridging the tendon ends in S100a4^GFP/+^ repairs compared to WT littermates (n=3-4 per group) (*) indicate sutures. (B-D) S100A4^GFP/+^ tendons expressed significantly more *Col1a1* mRNA (B), while transcript levels of *Col3a1* (C) and *Scx* (D) were unaffected by S100a4 haploinsufficiency. (*) indicates p<0.05 (un-paired t-test), n=3 per group. (E & F) *α-SMA* mRNA expression was significantly decreased in S100a4^GFP/+^ repairs (E) (n=3 per group), while a substantial reduction in α-SMA protein expression was observed in S100a4^GFP/+^, relative to WT, using immunofluorescence. White arrows indicate areas of α-SMA^+^ cells in the healing tissue, while yellow arrowheads denote α-SMA staining of vessels (F).

### S100a4 modulates macrophage function

Given the fibrotic nature of scar-mediated tendon healing, and the ability of macrophages to modulate multiple aspects of the fibrotic process(20–23), we examined changes in macrophage content during healing in S100a4 haploinsufficient mice, and changes in migration and polarization in response to S100a4 in vitro. A substantial decrease in F4/80^+^ macrophages were observed in S100a4^GFP/+^ repairs, relative to WT (white arrows, Fig. 4A), suggesting that knock-down of S100a4 can suppress macrophage recruitment or retention during tendon healing. S100a4-RP was shown to drive macrophage migration relative to vehicle, with a significant increase at the highest dose of 1000ng/mL (Fig. 4B). S100a4-RP treatment of primary macrophages appeared to have a variable impact on polarization. M1 markers *iNOS* and *CD64* were significantly upregulated in a dose-dependent manner following S100a4-RP treatment, while *TNFα* was downregulated and no change observed in *CD86* expression (p < 0.05) (Fig. 4C). M2 markers *Arg1* and *IL1ra* were significantly upregulated with higher doses of RP-S100a4, *CD163* was downregulated and no change seen in *CD206* (p<0.05) (Fig. 4D). Collectively this data supports a role for extracellular S100a4 in regulating macrophage migration and polarization.

**Figure 4.**
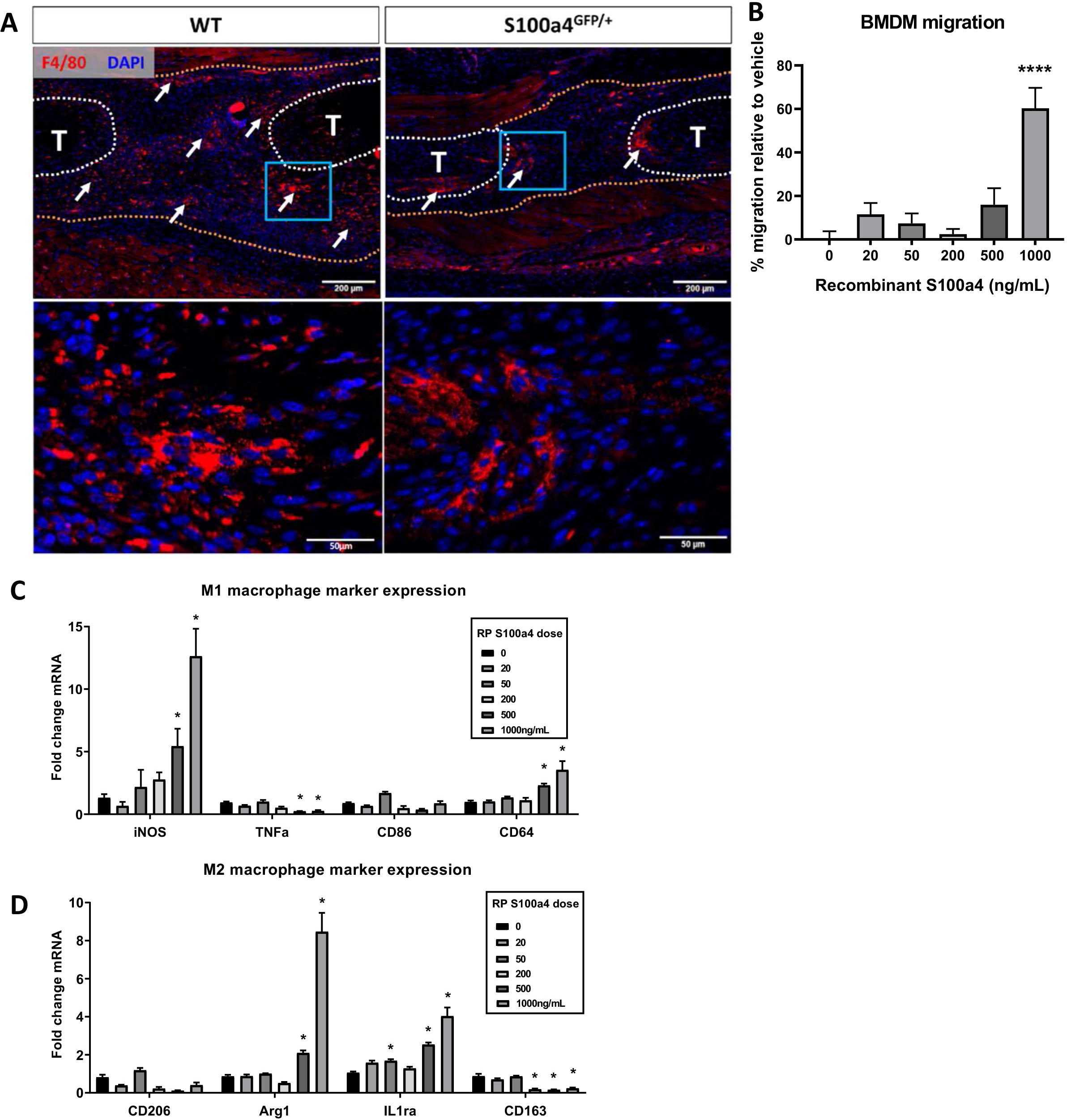
S100a4 alters macrophage function and response to tendon injury. (A) F4/80 staining demonstrates decreased macrophage content in the healing tendon of S100a4^GFP/+^ repairs. White arrows identify concentrated areas of macrophages. Tendon ends are outlined in white, while scar tissue is outlined in yellow (n=3-4 per group). (B) S100a4 promotes BMDM migration (n=3 per treatment). (****) Indicates p<0.0001 vs. vehicle treated cells (1-way ANOVA). (C & D) Following treatment with S100a4-RP (20-1000ng/mL), significant increases in M1 polarization markers (*iNos*, CD64) were observed relative to vehicle treated primary macrophages, a significant decrease in *TNFα*, and no change in *CD86* expression (C). Significant increases in M2 markers *Arg2* and *IL1ra* were seen with S100a4-RP treatment, a decrease in *CD163* expression, and no change in *CD206* (D) (n=3 per treatment). (*) indicates p<0.05 between vehicle and S100a4-RP treatment (1-way ANOVA).

### Inhibition of S100a4 signaling, via antagonism of RAGE improves tendon healing

Considering that knockdown of S100a4 expression improves healing, and S100a4 can function as an extracellular signaling molecule to drive fibrotic progression (24–26), we next examined expression of the putative S100a4 receptor, RAGE (Receptor for Advanced Glycation Endproducts) (27–29). RAGE expression was observed throughout the scar tissue during tendon healing, with abundant co-localization of S100a4 and RAGE (Fig. 4A). Consequently, we investigated the feasibility of inhibiting S100a4 signaling via disruption of S100a4-RAGE interaction using RAGE antagonist peptide (RAP)(30). In vivo, RAP treatment (Fig. 5B) significantly improved measures of gliding function relative to vehicle treated controls, with a 41% increase in MTP Flexion Angle (p=0.008) (Fig. 5C), and a 39% decrease in Gliding Resistance (p=0.007) (Fig. 5D). No differences in maximum load at failure (p=0.57) and stiffness (p=0.30) were observed between groups (Fig. 5E, F). These data suggest that inhibition of S100a4-RAGE recapitulates the improvements in gliding function seen with S100a4 haploinsufficiency but is insufficient to improve mechanical properties.

**Figure 5.**
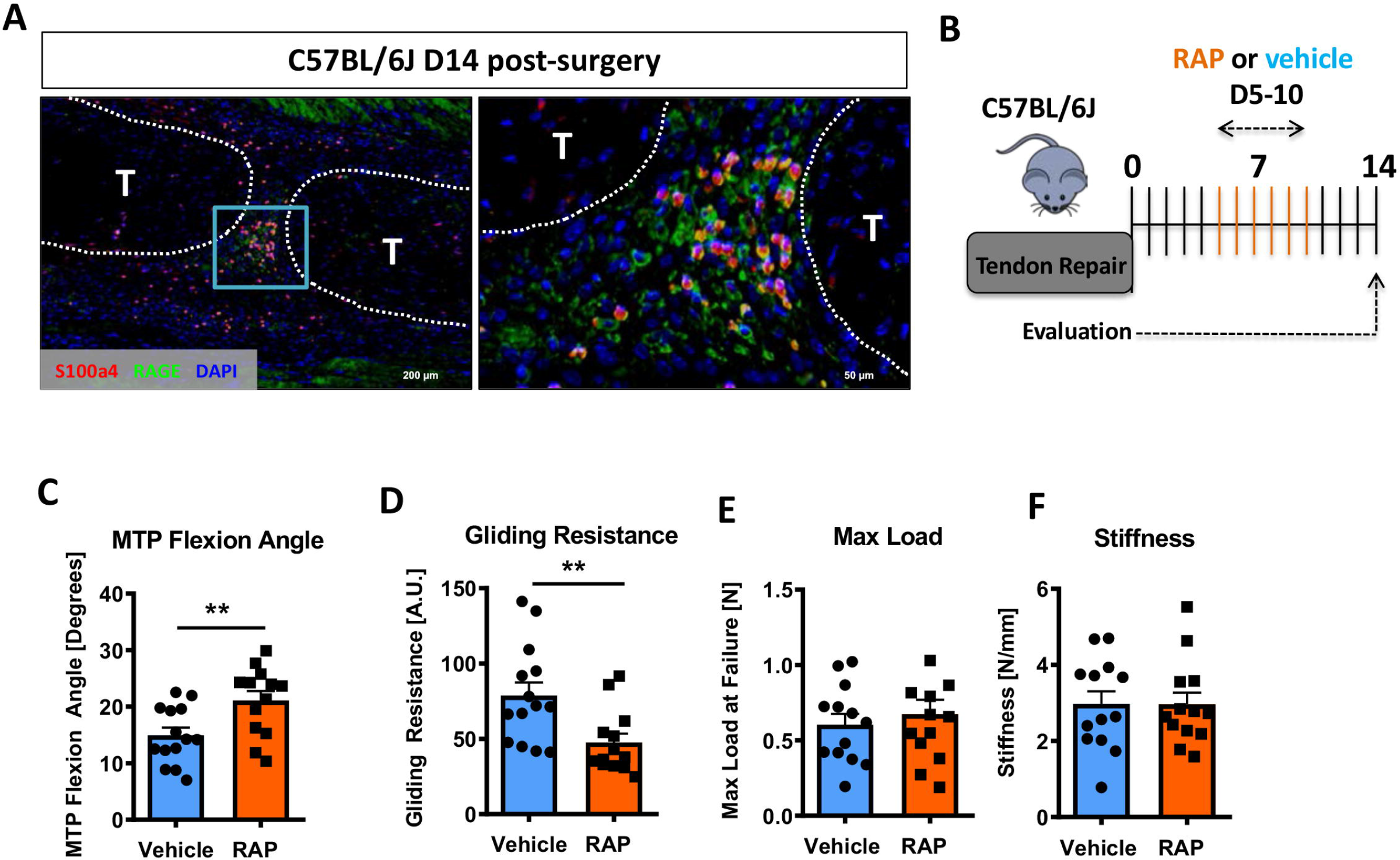
Inhibition of S100a4 signaling via RAGE antagonism improves tendon healing. (A) Co-immunofluorescence demonstrated co-localization of S100a4 and its putative receptor RAGE in the healing tendon (n=3). (B) C57Bl/6J mice were treated with either RAP or vehicle, via i.p. injection from D5-10 post-surgery, and harvested at D14 for functional testing. (C-F) At D14 RAP treatment significantly improved measures of gliding function relative to vehicle, with a (C) significant increase in MTP Flexion Angle, and (D) a significant decrease in Gliding Resistance. No change in (E) Max load at failure, or (F) Stiffness was observed between treatments (n=13 per group). (**) indicates p<0.01 between treatments (un-paired t-test).

### S100a4^+^ cell ablation results in aberrant matrix deposition during tendon healing

While knock-down of S100a4 expression and inhibition of S100a4 signaling improves tendon healing, S100a4 can also function in a cell-autonomous manner (31). To determine the effects of S100a4^+^ cell ablation on tendon healing, we depleted proliferating S100a4^+^ cells from D5-10 post-surgery (immediately preceding peak *S100a4* expression) (Fig. 6A). Depletion of S100a4^+^ cells from D5-10 resulted in a significant 91% reduction in *S100a4* mRNA expression (p<0.001) (Fig. 6B), and a substantial reduction in S100a4 protein expression, relative to WT (Fig. 6C). Functionally, slight but non-significant improvements in gliding function were observed in S100a4-TK (D5-10), relative to WT (Fig. 6D & E). However, max load at failure was significantly decreased by 43% (p=0.02) (Fig. 6F), while stiffness was unchanged (Fig. 6G). Morphologically, S100a4-TK (D5-10) tendons healed with thinner, more acellular bridging scar tissue between the native tendon ends, compared to the larger, more cellular granulation tissue in WT repairs (Figure 6H). Picrosirius staining demonstrated a substantial reduction in bridging ECM in S100a4-TK (D5-10) repairs, relative to WT (Fig. 6I). In contrast to this, qPCR revealed significant increases in ECM proteins *Col1a1* (3.9-fold, p=0.04) (Supplemental Figure 3A.) and *Col3a1* (1.9-fold, p=0.033) (Supplemental Figure 3B) mRNA expression in S100a4-TK (D5-10) mice. Additionally, significant decreases in the tenogenic transcription factor *Scx* (1.75-fold, p=0.04), and the myofibroblast marker α-SMA (9-fold, p=0.0003) were observed in S100a4-TK (D5-10), relative to WT (Supplemental Figure 3C & D). Consistent with this, and the phenotype in S100a4^GFP/+^ repairs, α-SMA staining was markedly reduced in S100a4-TK (D5-10) repairs, relative to WT (Supplemental Figure 4), as was total macrophage content (Supplemental Figure 5). Taken together, S100a4-cell depletion alters normal matrix deposition during tendon healing, leading to pronounced morphological changes in the scar tissue and a loss of overall strength, while recapitulating the changes in myofibroblast and macrophage populations observed in S100a4^GFP/+^ repairs.

**Figure 6.**
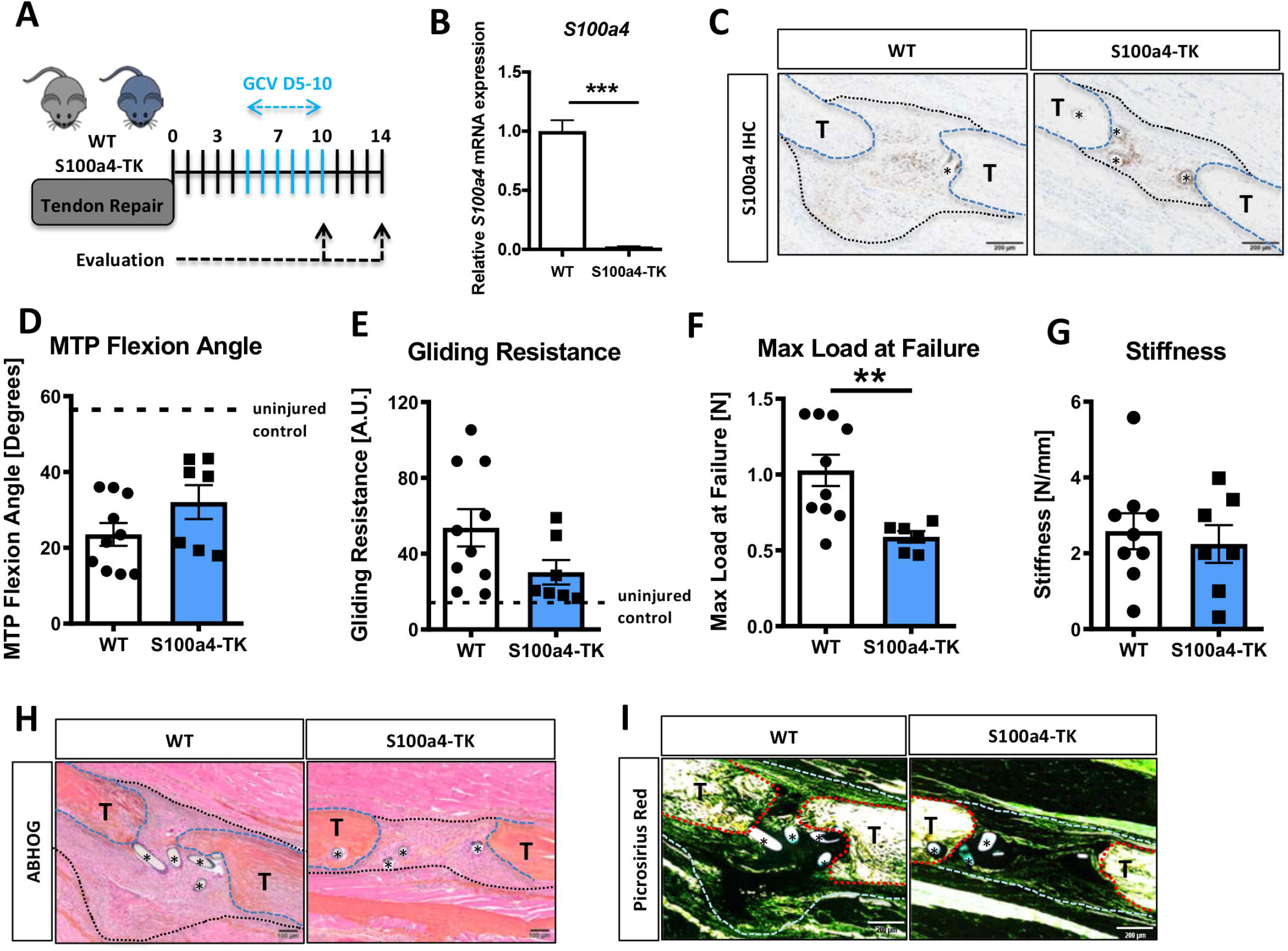
Delayed depletion of S100a4^+^ cells impairs restoration of mechanical properties and alters matrix deposition. (A) WT and S100a4-TK mice were treated twice daily with ganciclovir (GCV) from D5-10 post-surgery. (B) S100a4^+^ cell depletion results in a 91% reduction in S100a4 mRNA at D10 post-surgery (n=3). (C) A substantial reduction in S100a4 protein expression was observed S100a4-TK repairs, relative to WT. Tendon is outlined in blue, scar tissue is outlined in black and (*) identify sutures (n=4). (D-G) At D14 no change in MTP Flexion Angle (D) and Gliding Resistance (E) were observed between WT and S100a4-TK repairs. (F) Max load at failure was significantly reduced following S100a4-cell depletion, while no change in Stiffness was observed (G) (n=7-10), (**) indicates p<0.01 (un-paired t-test). (H & I) Morphologically, (H) ABHOG and (I) Picrosirius staining demonstrate reduced matrix deposition bridging the tendon ends in the S100a4-TK repairs, relative to WT. (*) Indicates sutures.

We then investigated the effect of continuous S100a4 cell depletion (D1-14) on healing (Figure 7A). In contrast to depletion from D5-10, depletion from D1-14 significantly reduced MTP Flexion Angle (−53%, p=0.0003) (Fig. 7B), and increased gliding resistance (+187%, p<0.001) (Fig. 7C), indicating impairment of normal gliding function with sustained S100a4^+^ cell depletion. Consistent with D5-10 depletion, depletion from D1-14 reduced mechanical properties, with a 43% decrease in max load (p=0.025) (Fig. 7D), and a 49% decrease in stiffness (p=0.0078) (Fig. 7E).

**Figure 7.**
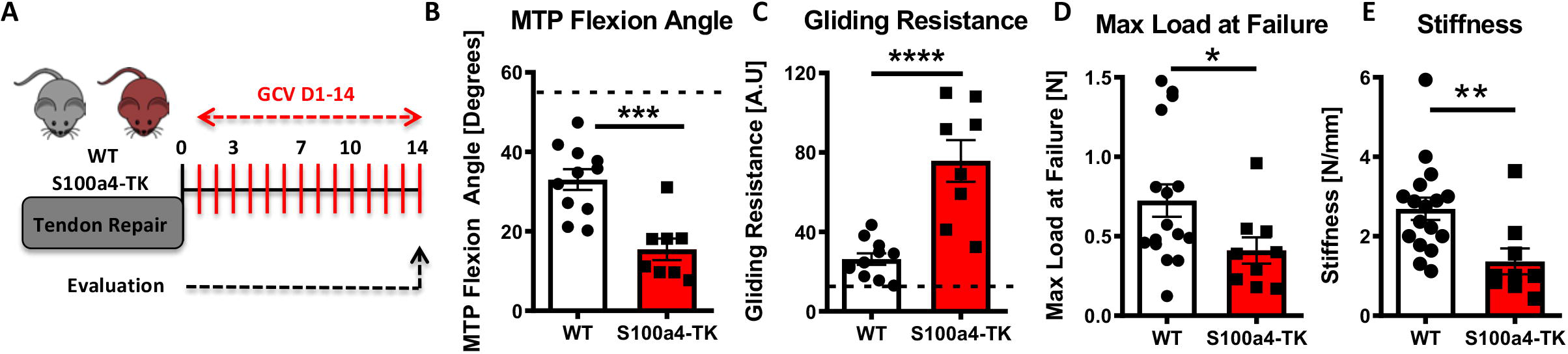
Sustained ablation of S100a4^+^ impairs restoration of gliding function and mechanical properties. (A) WT and S100a4-TK mice were treated with GCV from D1-14 post-surgery to ablate proliferating S100a4^+^ cells. At D14 (B) MTP Flexion Angle was significantly reduced, and (C) Gliding Resistance was significantly increased in S100a4-TK repairs, relative to WT. (D) A non-significant decrease in Max load at failure and (E) a significant reduction in Stiffness were observed in S100a4-TK repairs (n=8-11). (*) indicates p<0.05, (**) indicates p<0.01 (un-paired t-test).

### S100a4-lineage cells represent differentiated a-SMA myofibroblasts in the scar tissue of healing tendon

Examination of S100a4^GFP/+^ and S100a4-TK (D5-10) healing tendons demonstrate dramatically reduced myofibroblast content, suggesting potential interplay between these populations. The relationship between S100a4 and pro-fibrotic myofibroblasts is controversial and likely tissue-dependent, with conflicting reports of myofibroblast fate for S100a4^+^ cells(32–35). To understand the relationship between these cell populations during tendon healing we examined α-SMA expression in both S100a4-lineage cells and cells actively expressing S100a4. S100a4-Cre; Ai9 mice demonstrate that most α-SMA^+^ myofibroblasts at D14 post-surgery are derived from S100a4-lineage, as shown by co-localization (Fig. 8A, arrows). In contrast, very few S100a4^active+^ cells demonstrated co-localization with α-SMA (Fig. 8B). Taken together these data suggest that S100a4-lineage cells lose S100a4 expression during the transition to α-SMA^+^ myofibroblasts.

**Figure 8.**
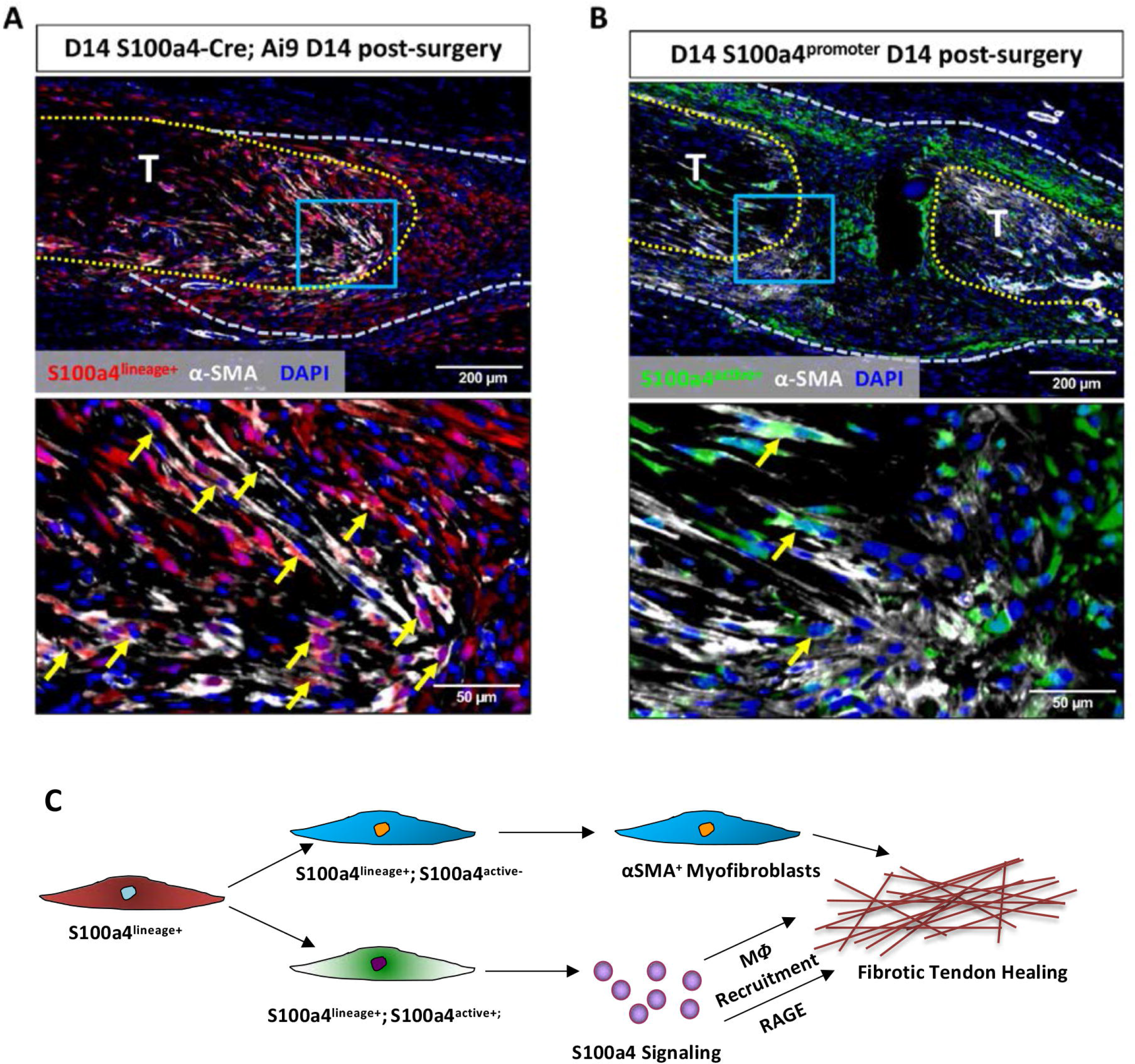
S100a4-lineage cells lose S100a4 expression during the transition to α-SMA^+^ myofibroblasts. (A) Co-localization of Red Fluorescent Protein (S100a4^lin+^ cells; red) and the myofibroblast marker α-SMA (white) demonstrated abundant co-localization (yellow arrows) during tendon healing. (B) Minimal co-localization of α-SMA (white) and cells actively expressing S100a4 (S100a4^active+^; green) was observed during healing (n=3). (C) Schematic representation of the cell non-autonomous signaling functions of S100a4 in fibrotic healing, as well as cell fate of S100a4-lineage cells.

## Discussion

Consistent with the role of S100a4 as a key driver of fibrosis (5, 14, 26, 36), we have demonstrated that S100a4 promotes scar-mediated tendon healing via cell-non-autonomous extracellular signaling. Further, we have shown that knockdown of S100a4 expression drives regenerative tendon healing and that depletion of S100a4^+^ cells disrupts re-acquisition of mechanical properties. Interestingly, the effects of S100a4^+^ cell depletion on functional metrics were time-dependent, suggesting a period of optimal S100a4 inhibition. Mechanistically, S100a4 interacts with RAGE and modulates macrophage content and function, suggesting the ability of S100a4 to regulate the inflammatory milieu during tendon healing. In addition, S100a4-lineage cells lose expression of S100a4 to become α-SMA^+^ pro-fibrotic myofibroblasts, which are likely involved in both restoration of matrix integrity and deposition of excess ECM. Taken together, these data establish S100a4 haploinsufficiency as a novel model of regenerative, mechanically superior tendon healing, and identify S100a4 as a potent anti-fibrotic therapeutic candidate to improve tendon healing.

S100a4 lacks enzymatic activity and functions predominantly through the regulation and interaction with other proteins. While the intracellular functions of S100a4 are not well-characterized, the extracellular signaling functions of S100a4 include regulation of multiple cellular processes important in fibrosis including motility (37, 38), differentiation (39–41) and survival (39). S100a4 protein levels are strongly correlated with idiopathic pulmonary fibrosis(34), and S100a4 has been suggested as a potential fibrotic biomarker in the liver (4). Consistent with this, we see highest S100a4 expression immediately prior to the period of peak scar formation during tendon healing. The therapeutic potential of targeting S100a4 has been well established in the lung, with decreased fibrotic progression following treatment with an S100a4 neutralizing antibody (34), and pharmacological inhibition of S100a4 (42). Moreover, Tomcik *et al.*, demonstrated that deletion of S100a4 prevented bleomycin-induced skin fibrosis(26). Consistent with these studies we demonstrated the knock-down of S100a4 is sufficient to attenuate scar-mediated tendon healing and promote a more regenerative response. More specifically, S100a4^GFP/+^ repairs demonstrate suppression of multiple components of the fibrotic cascade including macrophage content, myofibroblasts, as well as an altered ECM balance toward a mature tendon composition (Col1a1> Col3a1).

Macrophages play an essential role in wound healing, and S100a4 is a potent chemokine and regulator of macrophage chemotaxis. Bone marrow-derived macrophages from S100a4^−/−^ mice display defects in chemotactic motility and impaired recruitment to the site of inflammation (43). While we have not assessed the effects of S100a4 knockdown in macrophages, we have demonstrated that S100a4 alters macrophage migration and polarization in vitro and have shown that macrophage content is markedly reduced in S100a4 haploinsufficient and S100a4^+^ cell-depleted repairs, suggesting a potential role for extracellular S100a4 in macrophage recruitment during tendon healing. Moreover, while RAGE can be expressed on a variety of cell types, macrophages have been shown to be the primary source of RAGE in the context of pathology (44–47). Therefore, in addition to decreasing the macrophages content, and thereby RAGE^+^ cells, S100a4^GFP/+^ may also suppress signaling via direct down-regulation of RAGE. RAGE expression is highly dependent on ligand concentration (48), and we have decreased *S100a4* expression by 50%, implying down-regulation of the entire S100a4-RAGE-macrophage axis in S100a4^GFP/+^ mice. Interestingly, although inhibition of S100a4-RAGE with RAP recapitulates the improvements in gliding function observed in S100a4^GFP/+^ repairs, it is insufficient to improve mechanical properties. This suggests that S100a4 may regulate restoration of mechanical properties through mechanisms independent of RAGE. In addition to RAGE, S100a4 can interact with EGFR (49), Heparan Sulfate Proteoglycans (50), TLR4 (51) and Annexin 2 (52) receptors. However, these S100a4-receptor interactions have been primarily observed in metastasis and neurological disorders. Understanding the potential RAGE independent mechanisms of S100a4 signaling will be an important aspect of future research.

S100a4^+^ cell depletion has shown great efficacy in managing fibrotic progression in the peritoneum (10), and kidney (9). In contrast, depletion of S100a4^+^ cells impairs tendon healing, particularly when S100a4^+^ cells are depleted from D1-14. These opposing effects may be due to fundamental differences in the functions of affected tissues, with tendon being a mechanical, load-bearing tissue. Interestingly, the more pronounced negative effect of cell depletion from D1-14, relative to D5-10 depletion may be explained by suppression of the acute inflammatory phase. Anti-inflammatory administration during the acute inflammatory phase of tendon healing is effective at reducing scar formation, but causes marked reductions in mechanical properties (53–55). Taken together, these data further support the importance of timing treatment to modulate fibrotic tendon healing, particularly as it relates to S100a4 inhibition.

One of the main controversies surrounding S100a4^+^ cells is their potential to become α-SMA^+^ myofibroblasts. Osterreicher *et al.*, demonstrated that neither actively expressing S100a4^+^ cells or S100a4^lin+^ cells express α-SMA during lung fibrosis (56), and additional work demonstrates that S100a4^+^ bone marrow cells do not express α-SMA (57). In contrast, dermal fibroblasts are positive for expression of both S100a4 and α-SMA (56) and Chen *et al.*, demonstrate that S100a4 treatment increases α-SMA expression, while α-SMA^+^ cells decrease with S100a4-cell depletion in a model of liver fibrosis (4). Using a similar combination of S100a4-lineage tracing and active S100a4 expression analyses as in (56), we demonstrate that S100a4^lin+^ cells become α-SMA^+^, and that the α-SMA^+^ population is largely negative for S100a4. These data not only define a terminal myofibroblast fate for many S100a4-lineage cells during tendon healing, but further support the concept of exquisite cell and tissue specificity of S100a4 signaling. In addition to terminal cell fate, the origin of S100a4^+^ cells are unclear. Osterreicher *et al.*, demonstrate that bone marrow derived cells (BMDCs) represent the main source of S100a4 during liver fibrosis (56), and S100a4^+^ BMDCs have been shown to migrate to the site of neointima formation following vein grafting (57). While we have not traced S100a4^+^ BMDCs in this study, we have previously shown specific recruitment of BMDCs to the healing tendon (58), and we also see S100a4^Lin+^ and S100a4^active+^ populations in the tendon without injury. Future studies will seek to delineate the relative contribution of S100a4^+^ cells from different origins to scar-mediated tendon healing.

While we clearly identify S100a4 haploinsufficiency as a model of regenerative tendon healing, there are several limitations that must be considered. First, we have not investigated whether this signaling paradigm is conserved in other tendons. However, we do observe S100a4^+^ cells in the Achilles tendon at homeostasis and expansion of this population following injury, and since scar-mediated healing is consistent between tendons(59), this reinforces the potential application of this approach to improve healing in multiple tendons. Second, the long-term effects of inhibiting S100a4-RAGE signaling are unknown, as we have only examined healing at D14. However, peak expression of S100a4 at D10 post-surgery suggests the prime effects of S100a4 may occur during the early inflammatory/matrix deposition phases of healing. Moreover, the time-dependent effects of S100a4-cell depletion suggest there is likely an optimal therapeutic window for S100a4 inhibition, which will be examined in future studies. Finally, we have not delineated between the effects of S100a4 expression in resident tendon cells relative to expression in extrinsic cells, and how the cell origin of S100a4 may dictate the effects on healing.

Restoring satisfactory function following tendon injury has remained an intractable clinical problem for decades (60). To our knowledge this is the first model of regenerative tendon healing in transgenic mice, defined by improvements in range of motion and mechanics. These data will inform future work to define the pathways down-stream of S100a4-RAGE, rigorously determine the time-dependent effects of S100a4 inhibition and identify the cues that drive the transition of S100a4-lineage cells to myofibroblasts. More directly however, these studies define the tremendous potential of inhibition of S100a4 signaling as a therapeutic approach to promote regenerative tendon healing.

## Acknowledgements

We would like to thank the Histology, Biochemistry and Molecular Imaging (HBMI) and the Biomechanics, Biomaterials and Multimodal Tissue Imaging (BBMTI) Cores for technical assistance. This work was supported in part by NIH/ NIAMS K01AR068386 and R01AR073169 (to AEL). The HBMI and BBMTI Cores are supported by NIH/ NIAMS P30AR069655.

## Supplemental Figures

**Supplemental Figure 1. S100a4^+^ cells are found in the healthy and healing Achilles tendon.** S100a4-GFP^Promoter+^ (S100a4^active+^) cells are observed in the native Achilles tendon, and a S100a4^+^ population persists following complete transection and repair of the Achilles tendon at D14 post-surgery.

**Supplemental Figure 2. S100a4^GFP/+^ knock-in mice permit tracing of S100a4 haploinsufficient cells.** To determine if S100a4 haploinsufficiency altered the S100a4^+^ population during healing, fluorescent imaging of uninjured and D14 repairs from S100a4^GFP/+^ and S100a4^+/+^ (WT) mice were analyzed. No GFP expression was observed in S100a4^+/+^ WT mice either at baseline or at D14 post-surgery. In contrast, knock-down of S100a4 does not alter the S100a4^GFP+^ resident tendon cell population, or the expansion of the S100a4^GFP+^ population at D14 post-surgery. Orange insets identify high-power magnification images.

**Supplemental Figure 3. S100a4^+^ cell depletion alters expression of matrix, tenogenic and myofibroblast-associated genes.** qPCR analyses demonstrated significant increases in (A) *Col1a1*, and (B) *Col3a1* expression, while (C) *Scx* and (D) *α-SMA* expression levels were significantly reduced in S100a4-TK, relative to WT. Data were normalized to expression in WT samples and the internal control β-actin. (*) Indicates p<0.05, (***) indicates p<0.001, (n=3 per group)(un-paired t-test).

**Supplemental Figure 4. S100a4^+^ cell depletion reduces α-SMA^+^ myofibroblast content during healing.** At D14 post-surgery, abundant α-SMA^+^ myofibroblasts (red) were observed in WT repairs. In contrast, α-SMA^+^ myofibroblast content was markedly reduced in S100a4-TK (D5-10) repairs. White arrows indicate areas of α-SMA^+^ cells in the healing tissue, while yellow arrowheads denote α-SMA staining of vessels. Tendons are outlined in white and scar tissue outlined in red.

**Supplemental Figure 5. S100a4^+^ cell depletion reduces macrophage content during healing.** At D14 post-surgery abundant F4/80^+^ macrophages (red) were observed in WT repairs. In contrast, F4/80^+^ macrophage content was markedly reduced in S100a4-TK (D5-10) repairs. White arrows identify concentrated areas of macrophages. Tendon ends are outlined in white, while scar tissue is outlined in red. (*) indicates sutures.

Author contributions
Study conception and design: JEA, AEL; Acquisition of data: JEA, VS, KTB, EK; Analysis and interpretation of data: JEA, AEL; Drafting of manuscript: JEA, AEL; Revision and approval of manuscript: JEA, VS, KTB, EK, AEL.

## References

1. Pennisi E. Tending tender tendons. Science. 2002;295(5557):1011.

2. Aydin A, Topalan M, Mezdegi A, Sezer I, Ozkan T, Erer M, et al. [Single-stage flexor tendoplasty in the treatment of flexor tendon injuries]. Acta Orthop Traumatol Turc. 2004;38(1):54–9.

3. Louka ML, and Ramzy MM. Involvement of fibroblast-specific protein 1 (S100A4) and matrix metalloproteinase-13 (MMP-13) in CCl4-induced reversible liver fibrosis. Gene. 2016;579(1):29–33.

4. Chen L, Li J, Zhang J, Dai C, Liu X, Wang J, et al. S100A4 promotes liver fibrosis via activation of hepatic stellate cells. J Hepatol. 2015;62(1):156–64.

5. Lawson WE, Polosukhin VV, Zoia O, Stathopoulos GT, Han W, Plieth D, et al. Characterization of fibroblast-specific protein 1 in pulmonary fibrosis. Am J Respir Crit Care Med. 2005;171(8):899–907.

6. Tamaki Y, Iwanaga Y, Niizuma S, Kawashima T, Kato T, Inuzuka Y, et al. Metastasis-associated protein, S100A4 mediates cardiac fibrosis potentially through the modulation of p53 in cardiac fibroblasts. J Mol Cell Cardiol. 2013;57:72–81.

7. Yu CC, Tsai CH, Hsu HI, and Chang YC. Elevation of S100A4 expression in buccal mucosal fibroblasts by arecoline: involvement in the pathogenesis of oral submucous fibrosis. PloS one. 2013;8(1):e55122.

8. Flier SN, Tanjore H, Kokkotou EG, Sugimoto H, Zeisberg M, and Kalluri R. Identification of epithelial to mesenchymal transition as a novel source of fibroblasts in intestinal fibrosis. J Biol Chem. 2010;285(26):20202–12.

9. Iwano M, Fischer A, Okada H, Plieth D, Xue C, Danoff TM, et al. Conditional abatement of tissue fibrosis using nucleoside analogs to selectively corrupt DNA replication in transgenic fibroblasts. Mol Ther. 2001;3(2):149–59.

10. Okada H, Inoue T, Kanno Y, Kobayashi T, Watanabe Y, Ban S, et al. Selective depletion of fibroblasts preserves morphology and the functional integrity of peritoneum in transgenic mice with peritoneal fibrosing syndrome. Kidney international. 2003;64(5):1722–32.

11. Schneider M, Hansen JL, and Sheikh SP. S100A4: a common mediator of epithelial-mesenchymal transition, fibrosis and regeneration in diseases? Journal of molecular medicine (Berlin, Germany). 2008;86(5):507–22.

12. Björk P, Källberg E, Wellmar U, Riva M, Olsson A, He Z, et al. Common Interactions between S100A4 and S100A9 Defined by a Novel Chemical Probe. PLOS ONE. 2013;8(5):e63012.

13. Ackerman JE, Best KT, O’Keefe RJ, and Loiselle AE. Deletion of EP4 in S100a4-lineage cells reduces scar tissue formation during early but not later stages of tendon healing. Sci Rep. 2017;7(1):8658.

14. Iwano M, Plieth D, Danoff TM, Xue C, Okada H, and Neilson EG. Evidence that fibroblasts derive from epithelium during tissue fibrosis. Journal of Clinical Investigation. 2002;110(3):341–50.

15. Xue C, Plieth D, Venkov C, Xu C, and Neilson EG. The gatekeeper effect of epithelial-mesenchymal transition regulates the frequency of breast cancer metastasis. Cancer Res. 2003;63(12):3386–94.

16. Ackerman JE, and Loiselle AE. Murine Flexor Tendon Injury and Repair Surgery. J Vis Exp. 2016(115).

17. Hasslund S, Jacobson JA, Dadali T, Basile P, Ulrich-Vinther M, Soballe K, et al. Adhesions in a murine flexor tendon graft model: Autograft versus allograft reconstruction. J Orthop Res. 2008;26(6):824–33.

18. Loiselle AE, Bragdon GA, Jacobson JA, Hasslund S, Cortes ZE, Schwarz EM, et al. Remodeling of murine intrasynovial tendon adhesions following injury: MMP and neotendon gene expression. J Orthop Res. 2009;27(6):833–40.

19. Weischenfeldt J, and Porse B. Bone Marrow-Derived Macrophages (BMM): Isolation and Applications. CSH Protoc. 2008;2008:pdb prot5080.

20. Gibbons MA, MacKinnon AC, Ramachandran P, Dhaliwal K, Duffin R, Phythian-Adams AT, et al. Ly6Chi monocytes direct alternatively activated profibrotic macrophage regulation of lung fibrosis. Am J Respir Crit Care Med. 2011;184(5):569–81.

21. Murray LA, Chen Q, Kramer MS, Hesson DP, Argentieri RL, Peng X, et al. TGF-beta driven lung fibrosis is macrophage dependent and blocked by Serum amyloid P. Int J Biochem Cell Biol. 2011;43(1):154–62.

22. Wynn TA, and Ramalingam TR. Mechanisms of fibrosis: therapeutic translation for fibrotic disease. Nat Med. 2012;18(7):1028–40.

23. Wynn TA, and Vannella KM. Macrophages in Tissue Repair, Regeneration, and Fibrosis. Immunity. 2016;44(3):450–62.

24. Yammani RR, Carlson CS, Bresnick AR, and Loeser RF. Increase in production of matrix metalloproteinase 13 by human articular chondrocytes due to stimulation with S100A4: Role of the receptor for advanced glycation end products. Arthritis & Rheumatism. 2006;54(9):2901–11.

25. Miranda KJ, Loeser RF, and Yammani RR. Sumoylation and nuclear translocation of S100A4 regulate IL-1beta-mediated production of matrix metalloproteinase-13. J Biol Chem. 2010;285(41):31517–24.

26. Tomcik M, Palumbo-Zerr K, Zerr P, Avouac J, Dees C, Sumova B, et al. S100A4 amplifies TGF-beta-induced fibroblast activation in systemic sclerosis. Ann Rheum Dis. 2015;74(9):1748–55.

27. Grotterod I, Maelandsmo GM, and Boye K. Signal transduction mechanisms involved in S100A4-induced activation of the transcription factor NF-kappaB. BMC cancer. 2010;10:241.

28. Sorci G, Riuzzi F, Giambanco I, and Donato R. RAGE in tissue homeostasis, repair and regeneration. Biochim Biophys Acta. 2013;1833(1):101–9.

29. Donato R, Cannon BR, Sorci G, Riuzzi F, Hsu K, Weber DJ, et al. Functions of S100 Proteins. Current molecular medicine. 2013;13(1):24–57.

30. Arumugam T, Ramachandran V, Gomez SB, Schmidt AM, and Logsdon CD. S100P-derived RAGE antagonistic peptide reduces tumor growth and metastasis. Clin Cancer Res. 2012;18(16):4356–64.

31. Chow KH, Park HJ, George J, Yamamoto K, Gallup AD, Graber JH, et al. S100A4 Is a Biomarker and Regulator of Glioma Stem Cells That Is Critical for Mesenchymal Transition in Glioblastoma. Cancer Res. 2017;77(19):5360–73.

32. Humphreys BD, Lin SL, Kobayashi A, Hudson TE, Nowlin BT, Bonventre JV, et al. Fate tracing reveals the pericyte and not epithelial origin of myofibroblasts in kidney fibrosis. Am J Pathol. 2010;176(1):85–97.

33. Picard N, Baum O, Vogetseder A, Kaissling B, and Le Hir M. Origin of renal myofibroblasts in the model of unilateral ureter obstruction in the rat. Histochem Cell Biol. 2008;130(1):141–55.

34. Li Y, Bao J, Bian Y, Erben U, Wang P, Song K, et al. S100A4(+) Macrophages Are Necessary for Pulmonary Fibrosis by Activating Lung Fibroblasts. Front Immunol. 2018;9:1776.

35. Tanjore H, Xu XC, Polosukhin VV, Degryse AL, Li B, Han W, et al. Contribution of epithelial-derived fibroblasts to bleomycin-induced lung fibrosis. Am J Respir Crit Care Med. 2009;180(7):657–65.

36. Bruneval P, Rossert J, and Bariety J. Renewal of FSP1: a marker of fibrogenesis on human renal biopsies. Kidney international. 2005;68(3):1366–7.

37. Belot N, Pochet R, Heizmann CW, Kiss R, and Decaestecker C. Extracellular S100A4 stimulates the migration rate of astrocytic tumor cells by modifying the organization of their actin cytoskeleton. Biochim Biophys Acta. 2002;1600(1-2):74–83.

38. Schmidt-Hansen B, Klingelhofer J, Grum-Schwensen B, Christensen A, Andresen S, Kruse C, et al. Functional significance of metastasis-inducing S100A4(Mts1) in tumor-stroma interplay. J Biol Chem. 2004;279(23):24498–504.

39. Schneider M, Kostin S, Strom CC, Aplin M, Lyngbaek S, Theilade J, et al. S100A4 is upregulated in injured myocardium and promotes growth and survival of cardiac myocytes. Cardiovasc Res. 2007;75(1):40–50.

40. Stary M, Schneider M, Sheikh SP, and Weitzer G. Parietal endoderm secreted S100A4 promotes early cardiomyogenesis in embryoid bodies. Biochem Biophys Res Commun. 2006;343(2):555–63.

41. Novitskaya V, Grigorian M, Kriajevska M, Tarabykina S, Bronstein I, Berezin V, et al. Oligomeric forms of the metastasis-related Mts1 (S100A4) protein stimulate neuronal differentiation in cultures of rat hippocampal neurons. J Biol Chem. 2000;275(52):41278–86.

42. Zhang W, Ohno S, Steer B, Klee S, Staab-Weijnitz CA, Wagner D, et al. S100a4 Is Secreted by Alternatively Activated Alveolar Macrophages and Promotes Activation of Lung Fibroblasts in Pulmonary Fibrosis. Front Immunol. 2018;9:1216.

43. Li ZH, Dulyaninova NG, House RP, Almo SC, and Bresnick AR. S100A4 regulates macrophage chemotaxis. Mol Biol Cell. 2010;21(15):2598–610.

44. Gaens KH, Goossens GH, Niessen PM, van Greevenbroek MM, van der Kallen CJ, Niessen HW, et al. Nepsilon-(carboxymethyl)lysine-receptor for advanced glycation end product axis is a key modulator of obesity-induced dysregulation of adipokine expression and insulin resistance. Arterioscler Thromb Vasc Biol. 2014;34(6):1199–208.

45. Song F, Hurtado del Pozo C, Rosario R, Zou YS, Ananthakrishnan R, Xu X, et al. RAGE regulates the metabolic and inflammatory response to high-fat feeding in mice. Diabetes. 2014;63(6):1948–65.

46. Su XD, Li SS, Tian YQ, Zhang ZY, Zhang GZ, and Wang LX. Elevated serum levels of advanced glycation end products and their monocyte receptors in patients with type 2 diabetes. Arch Med Res. 2011;42(7):596–601.

47. Sunahori K, Yamamura M, Yamana J, Takasugi K, Kawashima M, and Makino H. Increased expression of receptor for advanced glycation end products by synovial tissue macrophages in rheumatoid arthritis. Arthritis Rheum. 2006;54(1):97–104.

48. Bierhaus A, Humpert PM, Morcos M, Wendt T, Chavakis T, Arnold B, et al. Understanding RAGE, the receptor for advanced glycation end products. J Mol Med (Berl). 2005;83(11):876–86.

49. Klingelhofer J, Moller HD, Sumer EU, Berg CH, Poulsen M, Kiryushko D, et al. Epidermal growth factor receptor ligands as new extracellular targets for the metastasis-promoting S100A4 protein. FEBS J. 2009;276(20):5936–48.

50. Kiryushko D, Novitskaya V, Soroka V, Klingelhofer J, Lukanidin E, Berezin V, et al. Molecular mechanisms of Ca(2+) signaling in neurons induced by the S100A4 protein. Mol Cell Biol. 2006;26(9):3625–38.

51. Cerezo LA, Remakova M, Tomcik M, Gay S, Neidhart M, Lukanidin E, et al. The metastasis-associated protein S100A4 promotes the inflammatory response of mononuclear cells via the TLR4 signalling pathway in rheumatoid arthritis. Rheumatology (Oxford). 2014;53(8):1520–6.

52. Semov A, Moreno MJ, Onichtchenko A, Abulrob A, Ball M, Ekiel I, et al. Metastasis-associated protein S100A4 induces angiogenesis through interaction with Annexin II and accelerated plasmin formation. J Biol Chem. 2005;280(21):20833–41.

53. Hammerman M, Blomgran P, Ramstedt S, and Aspenberg P. COX-2 inhibition impairs mechanical stimulation of early tendon healing in rats by reducing the response to microdamage. J Appl Physiol (1985). 2015;119(5):534–40.

54. Dimmen S, Engebretsen L, Nordsletten L, and Madsen JE. Negative effects of parecoxib and indomethacin on tendon healing: an experimental study in rats. Knee Surg Sports Traumatol Arthrosc. 2009;17(7):835–9.

55. Connizzo BK, Yannascoli SM, Tucker JJ, Caro AC, Riggin CN, Mauck RL, et al. The detrimental effects of systemic Ibuprofen delivery on tendon healing are time-dependent. Clin Orthop Relat Res. 2014;472(8):2433–9.

56. Osterreicher CH, Penz-Osterreicher M, Grivennikov SI, Guma M, Koltsova EK, Datz C, et al. Fibroblast-specific protein 1 identifies an inflammatory subpopulation of macrophages in the liver. Proc Natl Acad Sci U S A. 2011;108(1):308–13.

57. Cheng J, Wang Y, Liang A, Jia L, and Du J. FSP-1 silencing in bone marrow cells suppresses neointima formation in vein graft. Circulation research. 2012;110(2):230–40.

58. Loiselle AE, Frisch BJ, Wolenski M, Jacobson JA, Calvi LM, Schwarz EM, et al. Bone marrow-derived matrix metalloproteinase-9 is associated with fibrous adhesion formation after murine flexor tendon injury. PloS one. 2012;7(7):e40602.

59. Shepherd JH, Legerlotz K, Demirci T, Klemt C, Riley GP, and Screen HR. Functionally distinct tendon fascicles exhibit different creep and stress relaxation behaviour. Proceedings of the Institution of Mechanical Engineers Part H, Journal of engineering in medicine. 2014;228(1):49–59.

60. Strickland JW. Development of flexor tendon surgery: twenty-five years of progress. J Hand Surg [Am]. 2000;25(2):214–35.

