## Supplementary figures and images for "Cell non-autonomous functions of S100a4 drive fibrotic tendon healing"

### Supplemental Fig. 1

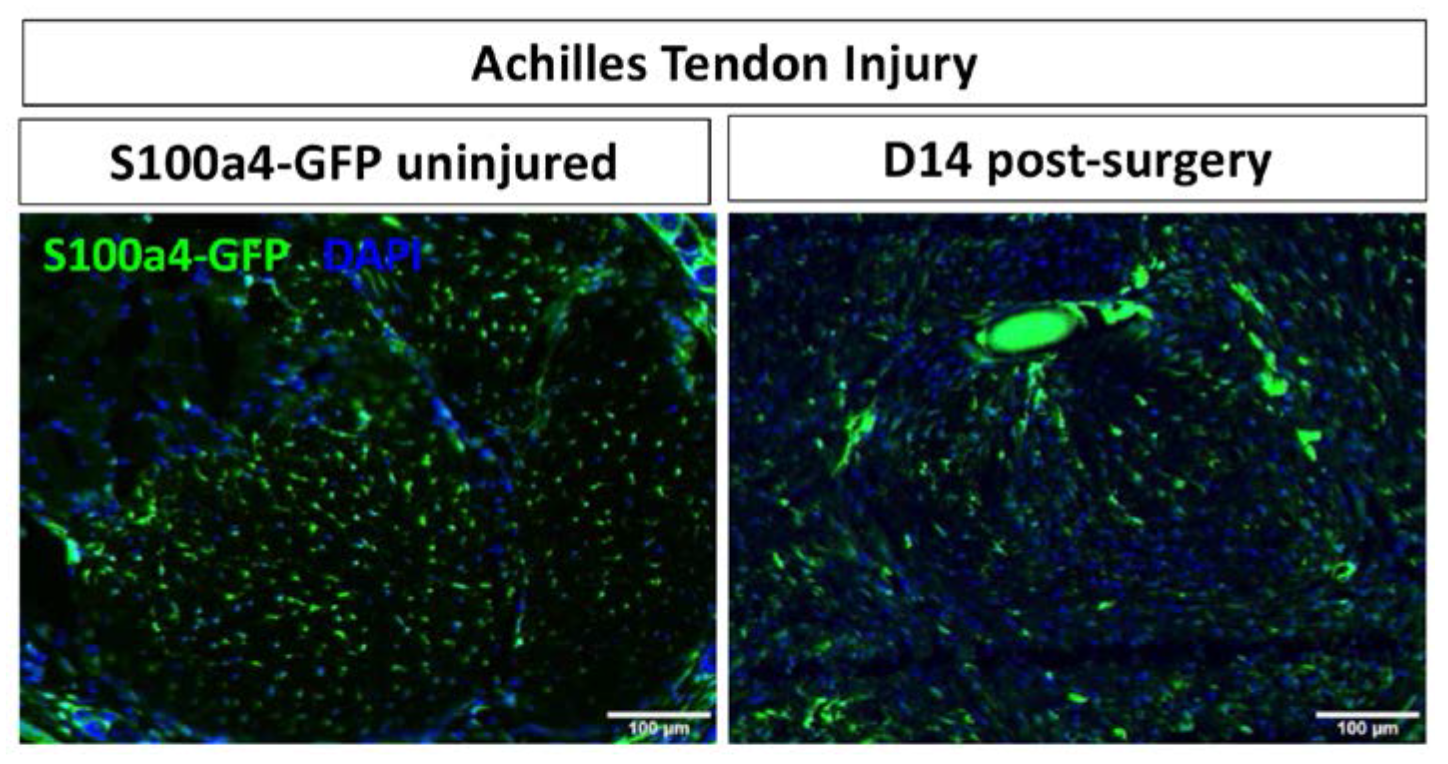

### Supplemental Fig. 2

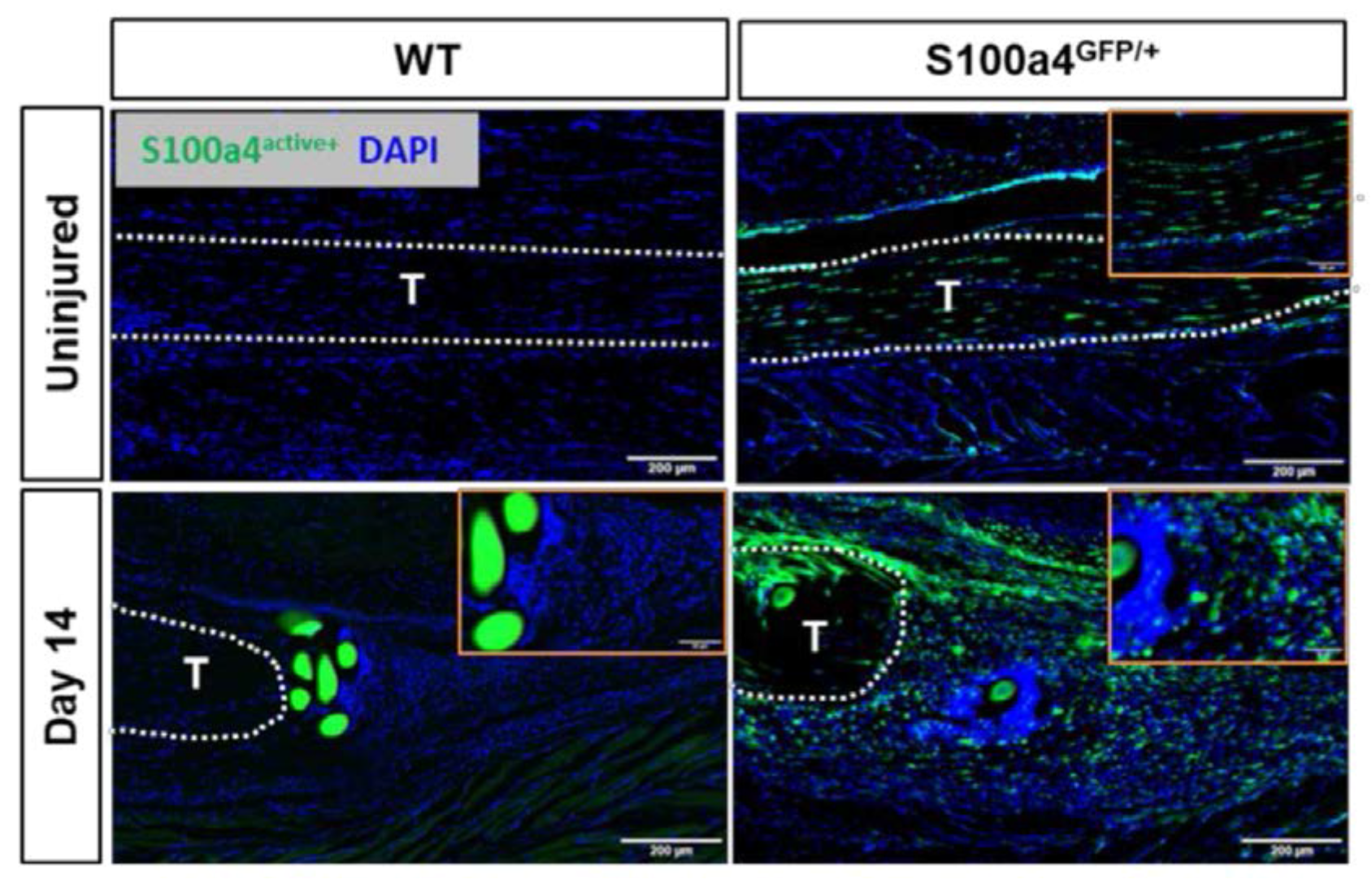

### Supplemental Fig. 3

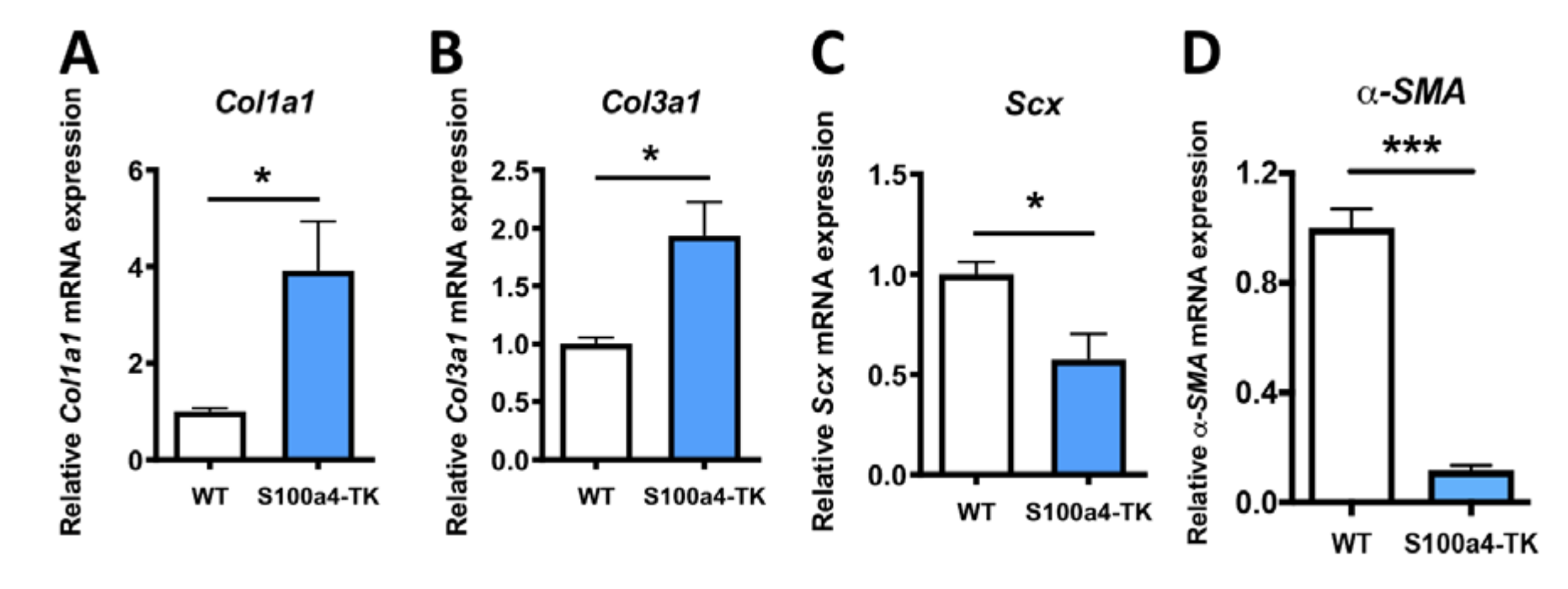

### Supplemental Fig. 4

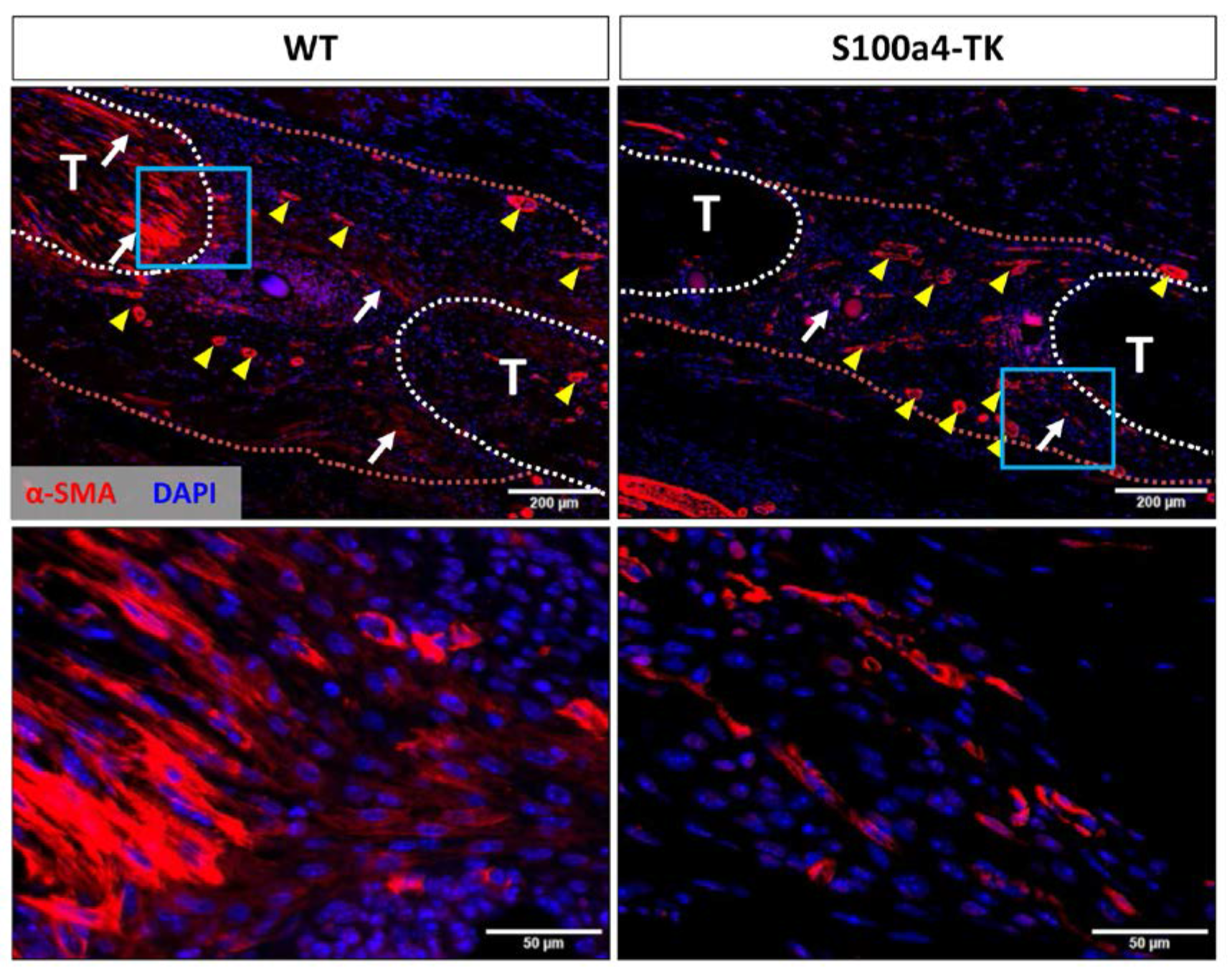

### Supplemental Fig. 5

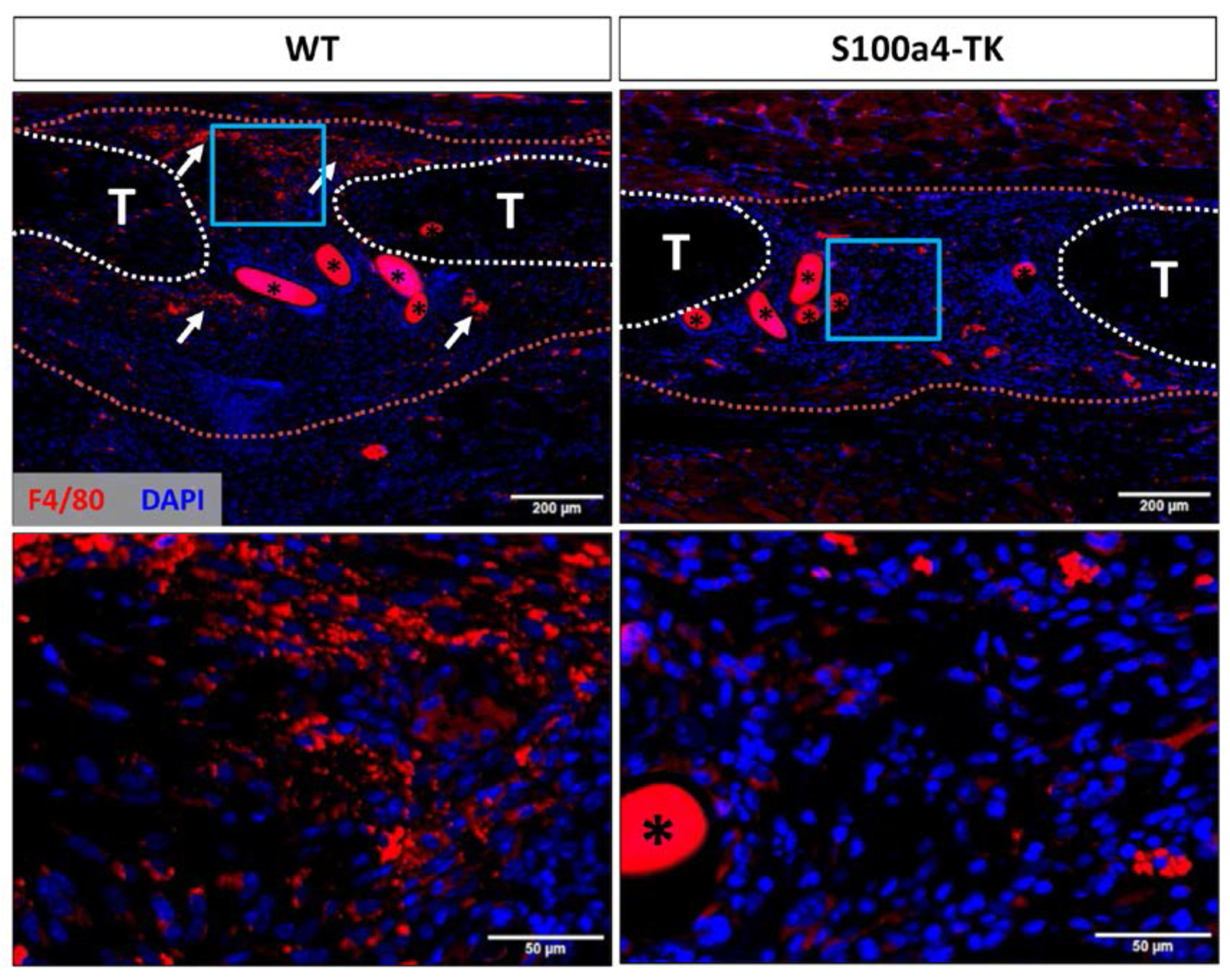
